# Remodeling hydrogen bond interactions results in relaxed specificity of Caspase-3

**DOI:** 10.1101/2020.08.07.241620

**Authors:** Liqi Yao, Paul Swartz, Paul Hamilton, A. Clay Clark

**Affiliations:** Department of Biology, University of Texas at Arlington, Arlington, Texas, 76019; Department of Molecular and Structural Biochemistry, NC State University, Raleigh, NC 27608; Department of Plant and Microbial Biology, North Carolina State University, Raleigh, NC 27695, USA

**Keywords:** enzyme specificity, apoptosis, caspase, zebrafish, protein evolution

## Abstract

Caspase enzymes play important roles in apoptosis and inflammation, and the non-identical but overlapping specificity profiles direct cells to different fates. Although all caspases prefer aspartate at the P1 position of the substrate, the caspase-6 subfamily shows preference for valine at the P4 position, while caspase-3 shows preference for aspartate. In comparison to human caspases, caspase-3a from zebrafish has relaxed specificity and demonstrates equal selection for either valine or aspartate at the P4 position. In the context of the caspase-3 conformational landscape, we show that changes in hydrogen bonding near the S3 subsite affect selection of the P4 amino acid. Swapping specificity with caspase-6 requires accessing new conformational space, where each landscape results in optimal binding of DxxD (caspase-3) or VxxD (caspase-6) substrate and simultaneously disfavors binding of the other substrate. Within the context of the caspase-3 conformational landscape, substitutions near the active site result in nearly equal activity against DxxD and VxxD by disrupting a hydrogen bonding network in the substrate binding pocket. The converse substitutions in zebrafish caspase-3a result in increased selection for P4 aspartate over valine. Overall, the data show that evolutionary neofunctionalization resulting in a dual function protease, as in zebrafish caspase-3a, requires fewer amino acid substitutions compared to those required to access new conformational space for swapping substrate specificity, such as between caspases-3 and −6.

## Introduction

Caspases, or cysteinyl-aspartate specific proteases, are important for apoptosis and cell differentiation (1), but the threshold of activity that results in differentiation or apoptosis is still unknown. The complex signaling pathways of eumetazoans utilize multiple caspases in three subfamilies, two of which are involved in cell death, and the third subfamily is involved in inflammation (2, 3). Caspases recognize a tetrapeptide sequence, with the exception of caspase-2 which prefers a pentapeptide sequence (2, 4), and all caspases prefer aspartate at the P1 position of the substrate. The P2 amino acid is generally hydrophobic, and the P3 amino acid is generally acidic and binds on the surface of the active site. Caspase specificity is defined primarily by the amino acid in the P4 position, such that caspases can be further classified into three groups based on their substrate preferences (Figure 1A): Group I, including caspase-1, −4 and −5, prefers the tetrapeptide sequence WEHD. Group II, including caspases-3 and −7, prefers DExD. Group III, including caspases-6, and −8, shows preferences for (I/L/V)ExD (x represents a non-specific amino acid) (5). Although the structures of all caspases are nearly identical, it is not clear which amino acid substitutions among homologs result in selection of the P4 amino acid. What may seem to be subtle differences in enzyme activity result in non-identical, yet overlapping, substrate specificities in the cell (6), so the selection of the P4 amino acid is important in cellular function.

**Figure 1.**
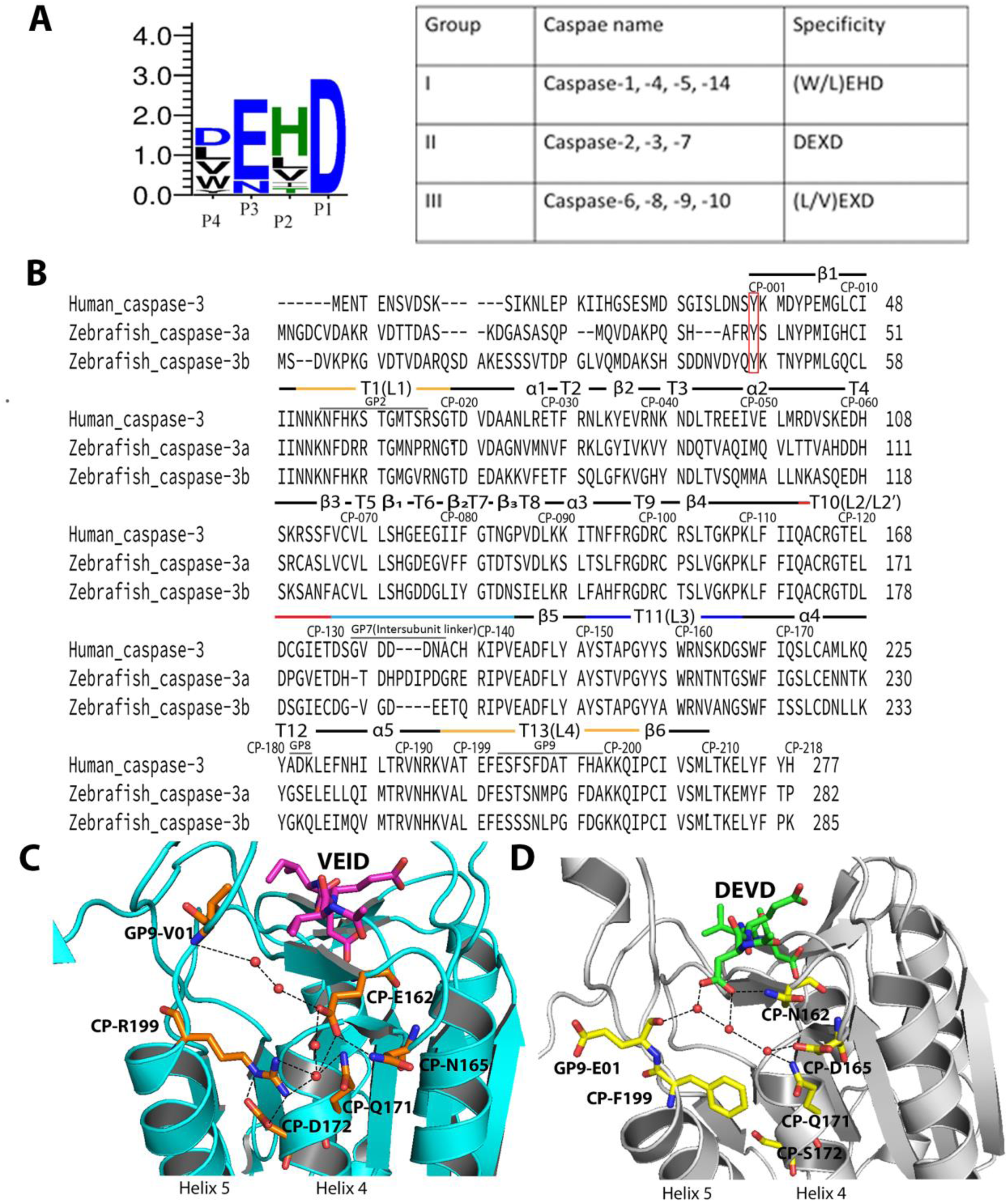
Properties of caspases. A. Combined sequence logo for substrate preferences in caspases (left) and substrate preferences for Group I-III caspases (right). B. Comparison of sequences for HsCasp-3, DrCasp-3a and DrCasp-3b. Common position numbering scheme and secondary structural elements are shown above each line, and actual position numbers are shown at the end of each row. Panels C-D. The charged amino acid network in HsCasp-6 (PDB ID 3S70) (panel C) is incomplete in HsCasp-3 (PDB ID 2J30) (panel D). In panels C and D, red spheres represent water molecules, and black dashed lines represent hydrogen bonds. Inhibitors are shown as purple (VEID, panel C) or green (DEVD, panel D).

Zebrafish (*Danio rerio*) caspases are excellent models for the study of protein evolution and enzyme specificity because zebrafish have 19 different caspase genes as a result of whole-genome duplication (7). Zebrafish have 2 copies of the caspase-3 gene, called DrCasp-3a and DrCasp-3b (Figure 1B). DrCasp-3a is expressed mainly in the brain and retina, while DrCasp-3b is expressed ubiquitously (7). In humans, HsCasp-3 is a DxxDase and acts as the primary executioner in apoptosis. It is expressed ubiquitously, while human caspase-6, a VxxDase, is expressed mainly in the brain or nervous system and controls cell development and differentiation (8). The localization of DrCasp3a and of DrCasp3b suggests that DrCasp-3a may be more similar to HsCasp-6 than to HsCasp-3, at least in terms of function during cell development. In support of this hypothesis, we showed previously that DrCasp-3a demonstrates equal activity against substrates with valine or aspartate at the P4 position (9).

Caspase subfamilies evolved from a common ancestor that diverged into multiple caspases more than 700 million years ago (10). We showed recently that the common ancestor of effector caspases-3, −6, and −7 was a promiscuous enzyme with little preference for aspartate or valine at the P4 site (11). In addition, specificity for valine evolved early in the caspase-6 lineage due to introduction of a network of charged amino acids near the S4 subsite (Figure 1C). The network results in a more hydrophobic S4 pocket by moving charged amino acids away from the pocket. It is not yet clear, however, how selection of aspartate evolved in the caspases-3,-7 lineages. The caspases-3,-7 proteins lack the charged amino acid network observed in caspase-6, which generally results in a more hydrophilic S4 pocket and hydrogen bonding interactions with the carboxylate at P4 (Figure 1D). In comparison, DrCasp-3a is similar in structure to HsCasp-3, in that it lacks the charged amino acid network observed in caspase-6 (9). Thus, the data suggest that within the framework of the caspase-3 conformational landscape, amino acid substitutions outside of the S4 subsite result in relaxed specificity of DrCasp-3a. Analysis of duplicated genes may show how genes diverged within the same genetic background, and in particular, how evolutionary changes in the caspase-3 conformational landscape affect enzyme selectivity.

Based on a comparison of HsCasp-3 and DrCasp-3a structures (9, 12), we generated a series of mutants to examine active site residues that affect substrate selection. We show that improving specificity for a P4 residue correlates to improved K_M_. For caspases, the K_M_ is thought to closely estimate KD, so the data suggest that selection correlates with improved binding of the P4 residue (2). For DrCasp-3a, with relaxed specificity, two amino acid substitutions near the S3 subsite shifted selection toward aspartate over valine, resulting from improved hydrogen bonding with the P3 residue. The converse substitutions in HsCasp-3 resulted in relaxed specificity and nearly equal activity against VxxD and DxxD. Thus, improved hydrogen bonding in the S3 subsite favors binding of aspartate in the P4 subsite while simultaneously disfavoring binding of the hydrophobic valine in the P4 subsite.

## Results and Discussion

### Comparing caspases from human and zebrafish suggest sites that affect specificity

We previously developed the common position (CP) numbering scheme in order to describe amino acid positions in caspases from multiple species (13). In the CP system, the amino acid positions that are common for each caspase are preceded by “CP-”, while regions that diverge are preceded by “GP-”. The GP, or gap regions, generally vary in number of amino acids and typically comprise loops near the active site or the intersubunit linker. The CP numbering for DrCasp-3a and HsCasp-3 is shown in Figure1B. Here we refer to the common position for consistency, but we provide the actual position number as well.

We used data from the CaspBase (13) and compared sequences and structures of caspases-3, −6 and −7, that is, the effector caspase subfamily. First, the analysis showed that the charged network in caspase-6, resulting in selection of P4 valine, is incomplete in caspases-3 and −7. While CP-E162 in caspase-6 is moved away from the S4 binding pocket due to the charged network, CP-N162 in caspase-3 forms a hydrogen bond with the carboxylate of the P4 aspartate (Figure 2A). In addition, conservation maps of amino acids between caspases-3 and −6 from mammals (Figure 2B) and between caspases-3 from mammals *versus* fish (Figure 2C) show similar patterns. That is, the core of the proteins as well as the active sites are reasonably well-conserved compared to the surface helices, which show much lower conservation. As we and others showed previously, the helices are part of allosteric networks which may be fine-tuned for species-specific function (14–16). Thus, the lower conservation in the helices may reflect increased evolutionary pressure on allosteric regulation of the enzyme, compared to the more conserved active site. More specifically, however, the comparison of the caspase-3, −6, −7 subfamily showed several amino acids in the active site with low conservation, even though the active site overall is more conserved. The amino acids are listed in Table 1 and include the charged amino acid network near the S4 binding pocket, described above, as well as two amino acids that are near a highly conserved arginine (CP-R161) that forms several hydrogen bonds with the P1 and P3 amino acids of the substrate (Figure 2D).

**Table 1.**
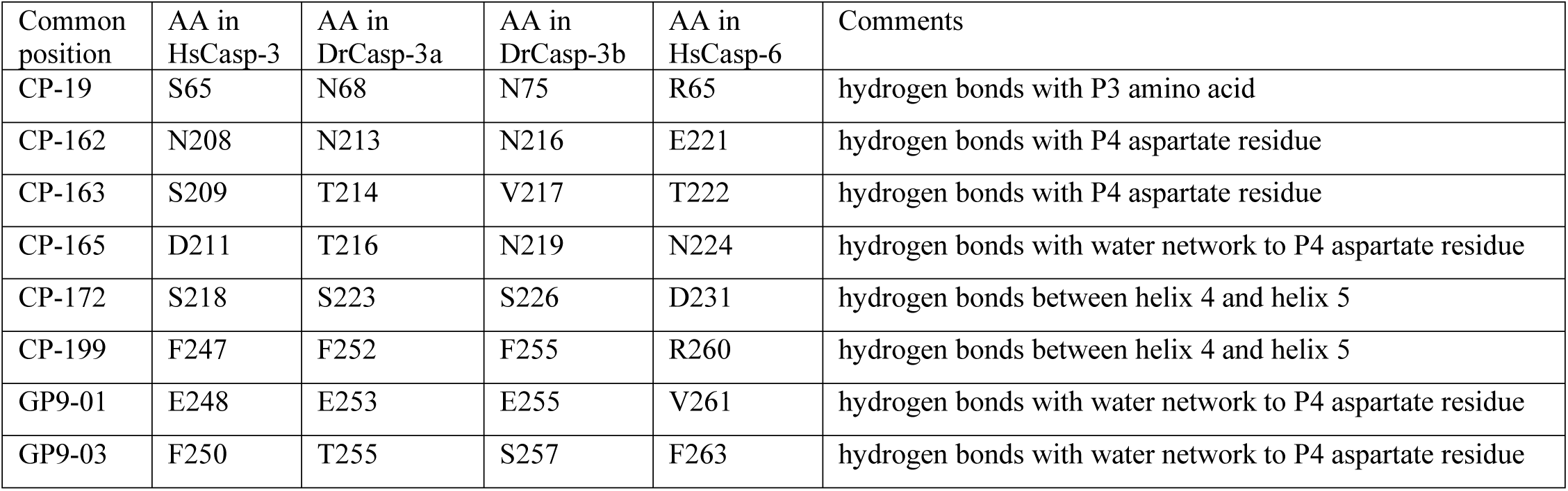
Comparison of amino acid positions that may affect specificity among human caspase-3, DrCasp-3a, and DrCasp–3b.

**Fig 2.**
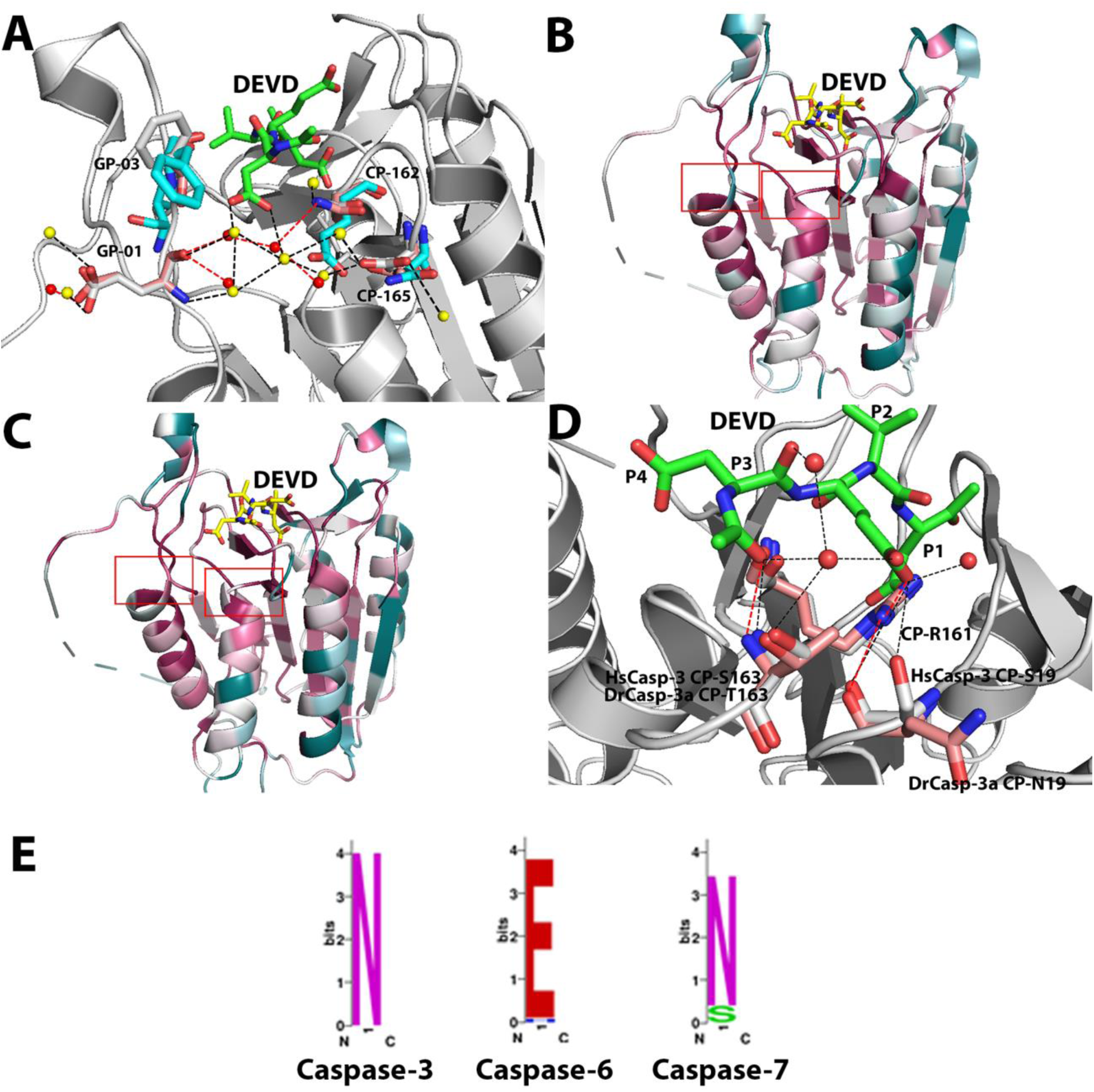
Comparison of human zebrafish caspases. A. Comparison of HsCasp-3 (grey), DrCasp-3a (salmon) and HsCasp-6 (cyan), showing CP-N162, CP-D165 and GP9-E01. The yellow spheres present the water molecules in HsCasp-3. The red spheres present water molecules in DrCasp-3a. The black dashed lines represent hydrogen bonds in HsCasp-3, while the red dashed lines represent hydrogen bonds in DrCasp-3a. B. Conservation map of comparison between caspase-3 and caspase-6 from mammals. C. Conservation map of comparison between caspase-3 from mammals and from fish. For panels B and C, sites with higher conservation are shown in red, and less conserved sites are shown in cyan. Conservation maps were determined using the ConSurf server (30–32). D. Comparison of HsCasp-3 (grey) and DrCasp-3a (salmon), showing CP-19, CP-212, and CP-163. Red spheres represent water molecules, and black dashed lines represent hydrogen bonds in HsCasp-3. E. Amino acids utilized at CP-162, based on multiple sequence alignments of caspase-3, −6 and −7 from 200 species. Asparagine is conserved in caspase-3 and −7, while glutamate is conserved in the caspase-6 lineage.

To determine the role, if any, of the amino acids listed in Table 1 in substrate selection, we first examined amino acids in or near the S4 subsite. As described previously (13), CP162 is conserved in caspase-6 (VxxDases) as glutamate (Figure 1, panels B, C, and Figure 2E). In contrast, CP162 is conserved in caspases-3 and −7 (DxxDases) as asparagine (Figure 1, panels B, D and Figure 2E). Reconstructions of ancestral effector caspases showed that the polar glutamate in the caspase-6 lineage is moved away from the S4 subsite by hydrogen bonding to a network of amino acids on helices-4 and −5, as described above (11). The interactions result in rotation of a hydrophobic group on active site loop 4 (L4), called GP9-V01, into the S4 subsite (Figure 1C) (11). The charged network on helices 4 and 5, observed in caspase-6, is incomplete in caspases-3 and −7, and the hydrophobic GP9-V01 is substituted with glutamate in the DxxDases (Figure 1D). Thus, within the framework of the conformational landscape for each caspase, the structural comparisons suggest that completing the charged amino acid network in helices 4 and 5, and substituting GP9-01E for valine, should shift specificity in caspase-3 toward P4 valine. To test the hypothesis, we introduced mutations in HsCasp-3 that would complete the charged network observed in caspase-6 (CP-N162E, CP-D165N, CP-S172D, CP-F199R, GP9-E01V; see Table 1). The data for the HsCasp-3 mutant, however, show that the mutant has a substantial increase in K_M_ for DEVD substrate and no activity toward VEID substrate (Table 2). In the context of the caspase-3 conformational landscape, simply completing the charged network observed in caspase-6 is not sufficient to change enzyme specificity.

**Table 2.**
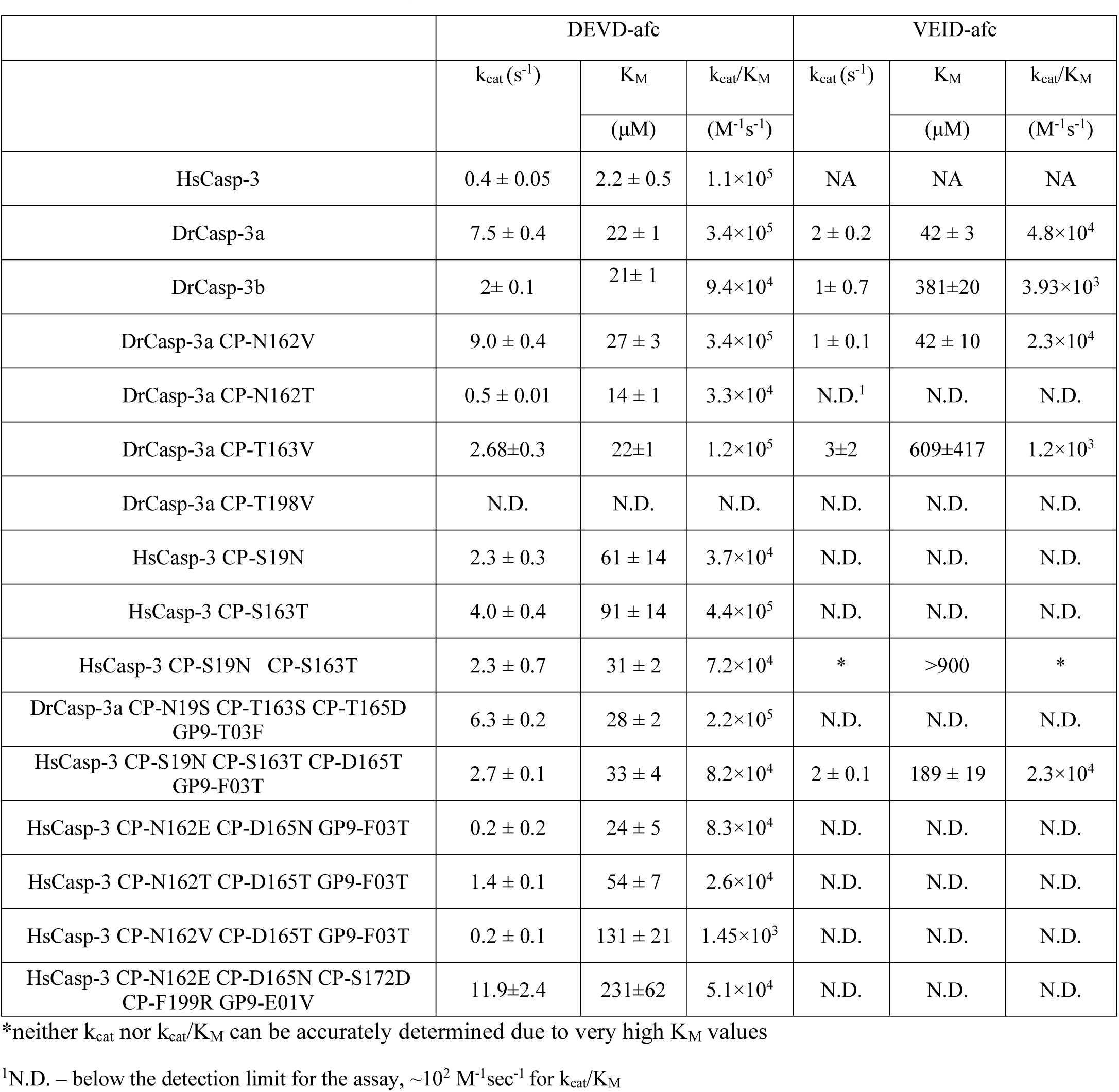
Catalytic parameters for caspase-3 and all mutants

Since the caspase-3 and −7 subfamilies do not contain the charged amino acid network observed in caspase-6, other evolutionary changes must also affect substrate selection within the framework of the caspase-3 conformational landscape for DrCasp-3a to exhibit similar activity against P4 aspartate or valine. In comparison to HsCasp-3 and HsCasp-6, DrCasp-3a is more structurally similar to HsCasp-3 in that CP-N162 and GP9-E01 are conserved, and the charged/polar network on helices 4 and 5 is incomplete (Table 1 and Figures 1B, 2A). Thus, it is not clear how changes in the conformational landscape of the DxxDase result in relaxed specificity to allow binding of aspartate or valine in the S4 subsite.

The whole-genome duplication in teleost fish resulted in two copies of caspase-3 in zebrafish, called DrCasp-3a and DrCasp-3b (7). Thus, one can examine neofunctionalization of the enzymes as they evolve together in the same organism. In the absence of structural data for DrCasp-3b, we generated a structural model based on our previous X-ray crystal structure of DrCasp-3a (9). The two enzymes have 60.6% identity at the amino acid level, and the model shows a similar structure to DrCasp-3a. Importantly, the S4 binding pocket is essentially identical to that of DrCasp-3a, with CP-N162 and GP9-E01 being conserved, and the polar network on helices 4 and 5 is incomplete, as observed for the DxxDases. As described below, DrCasp-3b is most selective for P4 aspartate and shows much lower activity against P4 valine. In addition, conservation maps, described above, between caspases-3 and −6 from mammals (Figure 2B) and caspases-3 from mammals *versus* fish (Figure 2C) show similar patterns overall, but more specifically, HsCasp-3 and DrCasp-3a sequences show 80 substitutions between the two proteins (Figure 1B), with 8 substitutions near the active site. In comparison, there are 75 differences between DrCasp-3a and DrCasp-3b, with 6 substitutions near the S4 and S3 binding pockets. Based on comparisons of the three proteins, we examined substitutions at eight sites, as shown in Table 1, including the five amino acids described above that comprise the charged network from caspase-6. Our goal was to determine interactions that result in the relaxed specificity for DrCasp-3a and that, in turn, result in the selection of aspartate over valine in HsCasp-3.

We first examined changes at CP-N162 by mutating the residue to Val or Thr in DrCasp-3a or to Val, Thr, or Glu in HsCasp-3. The rationale was to determine whether removing the hydrogen bond to the P4 aspartate and increasing hydrophobicity in the S4 subsite would result in changes in substrate selection. The results show little change in DrCasp-3a with CP-N162 to Val substitution, but the CP-N162 to Thr variant of DrCasp-3a demonstrated no activity against VEID substrate (Table 2, Supplementary Figure 1). In contrast, the activity against DEVD was about four-fold lower than that of the wild-type enzyme, due to an increase in K_M_ (Table 2). We determined the X-ray crystal structure of DrCasp-3a(CP-N162T) variant at 2.7 Å resolution, and the data show little to no change in the active site (Supplementary Figure 2). The hydroxyl group of the Thr side-chain forms a hydrogen bond with its backbone amide group, which presumably stabilizes the turn in active site loop 3 (called L3). The methyl group of the Thr side-chain is oriented toward the acetyl group of the substrate, while the P4 aspartate forms three hydrogen bonds with groups in the L3 and L4 loops. Thus, it is not clear from the structure why this variant has no activity against VEID substrate and a lower catalytic efficiency against DEVD substrate.

In HsCasp-3, we examined CP-N162 in the context of two additional sites (CP-D165 and GP9-F03). The latter two sites are either part of the charged network described above (CP-D165, in active site loop 3) or form a hydrophobic cluster on the surface of active site loop 4 (GP9-F03) (see Figure 2A). The substitution of CP-N162 for glutamate had little effect on activity against DEVD substrate, but substitutions of CP-N162 for threonine or valine had much larger effects, between 5- and 100-fold decrease in activity against DEVD substrate. Regardless of the substitution, however, the HsCasp-3 CP-N162 variants demonstrated no increase in activity against VEID (Table 2). Together, the structural models and activity data for the S4 subsite variants show that selection of valine or aspartate in the P4 position is affected by factors outside of the S4 subsite. That is, simply changing the hydrophobicity of the S4 subsite was not sufficient to change substrate selection. Overall, we interpret the results to show that, in the absence of other changes in the active site, the loss of the hydrogen bond between the P4 aspartate and CP-N162, the hydrophobic cluster on the surface of loop L4, and the hydrogen bonds contributed by CP-D165 are not critical factors for determining selection. Notably, the substitutions generally lower the activity against DEVD substrate without concomitant increase in activity against VEID.

We next examined two sites away from the S4 subsite (CP-19 and CP-163) due to differences among HsCasp-3, DrCasp-3a, and DrCasp-3b (Table 1 and Figure 2). In HsCasp-3, the side-chain of CP-S19 contributes to a hydrogen bonding network between several water molecules and the universally conserved CP-R161 (Figure 2D). The side-chain of CP-R161 is critical for substrate binding because it forms hydrogen bonds with the P3 and P1 side-chains as well as several residues in active site loops 1 and 3 (Figure 2D). In addition, several water molecules contribute to the hydrogen bonding network that coordinates the P1 and P3 residues of the substrate (Figure 2D). In DrCasp-3a and DrCasp-3b, CP-S19 is substituted with asparagine (Table 1, Figure 2D). Although CP-S19 directly hydrogen bonds to the P3 glutamate in HsCasp-3, the side-chain of CP-N19 is rotated toward solvent in DrCasp3a. A water molecule occupies the position of the hydroxyl group found in HsCasp-3, and the hydrogen bonding network is maintained through the water molecule and the backbone carbonyl of CP-N19 (Figure 2D). In contrast to CP-S/N19, the three proteins differ in the amino acid at CP-163, where HsCasp-3, DrCasp-3a, and DrCasp-3b utilize serine, threonine, or valine, respectively (Table 1). In HsCasp-3 and in DrCasp-3a, the hydroxyl group of the side-chain forms part of a water-mediated hydrogen bond with the P3 glutamate. In HsCasp-3, we first introduced substitutions of CP-S19N and CP-S163T in the context of the CP-D165T and GP9-F03T mutations described above. We also introduced comparable substitutions in DrCasp-3a (CP-N19S, CP-T163S, CP-T165D, GP9-T03F) (Table 2). The HsCasp-3 variant showed activity against DEVD similarly to that of wild-type HsCasp-3, with k_cat_/K_M_ values of 8.2×10^4^ and 1.1×10^5^ M^−1^sec^−1^, respectively, so the mutations had little effect on the DxxDase activity. However, the HsCasp-3 variant also demonstrated activity against VEID, albeit with K_M_ values ~5-fold higher than that of the wild-type DrCasp-3a (Table 2). Similarly, the DrCasp-3a variant that contained the comparable substitutions from HsCasp-3 showed high activity against DEVD, but little to no activity against VEID (Table 2 and Supplemental Figure 1). Based on our results described above for positions CP-D165 and GP9-F03, which demonstrated no change in substrate selection when combined with mutations at CP-N162, we reasoned that the results observed for mutants containing the four substitutions were due to the changes at CP-S19 or CP-S163, or both. We introduced single substitutions in HsCasp-3 to mimic DrCasp3a (CP-S19N, CP-S163T), and we also substituted both sites together in HsCasp-3. Both single variants showed activity against DEVD substrate, albeit with higher K_M_ values, and no activity against VEID substrate (Table 2 and Supplemental Figure 1). The double mutant also retained high activity against DEVD substrate, and there was observable activity against VEID substrate. However, a very high K_M_ for VEID prevented an accurate determination of k_cat_ as well as the specificity constant, k_cat_/K_M_ (Table 2).

We confirmed the changes in substrate selection for the caspase-3 variants using phage-display substrate assays (Supplemental Figure 3). In those assays, immobilized substrate phage libraries randomized from P5-P1’, or fixed with aspartate at P1, are cleaved with caspase variants, and the preferred sequences are determined after several rounds of selection and amplification (9). Whereas wild-type DrCasp-3a shows selection for valine or aspartate at P4, wild-type HsCasp-3 demonstrates selection for aspartate but not valine (9). Our results also show that DrCasp-3b exhibits a preference for P4 aspartate (Supplemental Figure 3A), which is consistent with peptide substrate studies (Table 2). In contrast to DrCasp-3a, which exhibits high activity against VEID substrate, DrCasp-3b shows low, but measurable, activity against P4 valine (Table 2). In this case, the K_M_ for VEID substrate is ~10-fold higher in DrCasp-3b compared to DrCasp3a. In addition, the HsCasp-3 variants that included substitutions at CP-S19 and CP-S163 exhibit lower selection for P4 aspartate and increased selection for valine, serine, or threonine at P4 (Supplemental Figure 3). Together, the data show that substitutions near the S1 and S3 pockets, CP-19 and CP-163, when combined with substitutions near the S4 subsite, CP-165 and GP9-03, result in relaxed specificity for HsCasp-3 as well as increased selection for P4 aspartate in DrCasp-3a.

We examined the effects of the substitutions at CP-19 and CP-163 by molecular dynamics simulations (50 ns) in order to examine changes in mobility of the active site (Figure 3). The results show that the mutations had little effect on DEVD substrate bound in the active site (Figure 3A). In the case of VEID substrate, however, the peptide at the P1 and P2 positions was released from the binding pocket (Figure 3B). The data show that in wild-type DrCasp-3a, the mobility of CP-R161 is restrained due to the hydrogen bonding network with CP-N19, CP-T163, and the network of water molecules (Figures 3A and 3C). This is true regardless of DEVD or VEID substrate bound in the active site. In contrast, substitutions at CP-19 and CP-163 resulted in increased mobility of CP-R161 with VEID substrate, but not DEVD substrate, bound in the active site (Figure 3D). In the case of the VEID substrate, the increased mobility of CP-R161 resulted in clashes with the peptide backbone near the P2 residue. The clashes likely result in increased mobility of the peptide at the P1 and P2 sites. We suggest that with DEVD bound, the increase in hydrogen bonding with the P4 aspartate provides sufficient stability to prevent the increased movement of CP-R161 in the active site. In contrast, the binding of valine in the S4 subsite, and lack of hydrogen bonding with the side-chain, is insufficient to prevent mobility of CP-R161 and subsequent steric clashes with the bound substrate.

**Figure 3.**
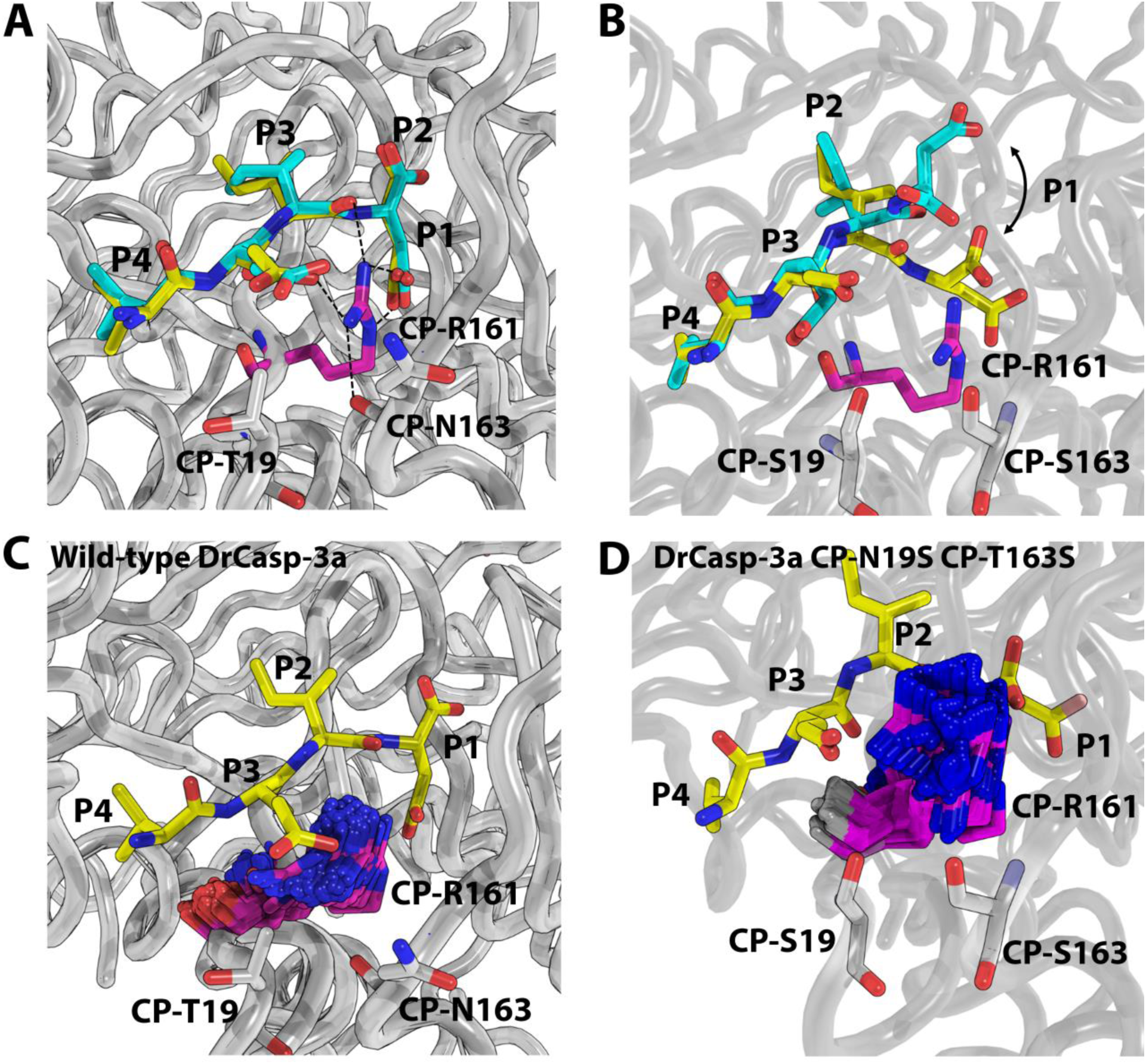
Molecular dynamics simulations for CP-19S/N and CP-163S/T. A. Wild-type DrCasp-3a with VEID bound at time zero (yellow) and at 17 ns (cyan). B. DrCasp3a(CP-N19S,CP-T163S) variant with VEID bound at time zero (yellow) and at 17 ns (cyan). Panels C,D. Comparison of mobility in CP-R161 in wild-type (panel C) versus CP-N19S,CP-T163S mutant of DrCasp3a. In both panels C and D, the position of VEID substrate is shown at the beginning of the simulation, and the position of CP-R161 is shown every 0.25 ns over the course of the 50 ns simulation.

## Conclusions

We have shown that hydrogen bonding is an important determinant for selection of the P4 residue of caspase substrates and that interactions outside of the S4 binding pocket contribute to the selection of the P4 residue. In the conformational landscape of caspase-6, a network of charged or polar amino acids moves CP-E162 away from the S4 pocket, and rotation of GP9-V01 into the pocket increases hydrophobicity and thus preference for P4 valine in the substrate (11). In the context of the caspase-3 conformational landscape, however, nature did not use this strategy to change selection toward P4 aspartate. In contrast to the single caspase-3 in humans, zebrafish have two caspase-3 enzymes due to a whole-genome duplication (7), and the two caspase-3 enzymes exhibit differences in substrate selection. We showed previously that DrCasp-3a exhibits relaxed specificity, with approximately equal activity against DEVD and VEID (9). Our data described here shows that DrCasp-3b is more similar to HsCasp-3, with preference for P4 aspartate. We show that two sites, on active site loops L1 and L3, when combined with two sites near the S4 binding pocket, contribute to a hydrogen bonding network that stabilizes the P1 and P3 residues of the DxxD substrate by decreasing the mobility of a conserved arginine residue that interacts with the side-chains of both amino acids. Thus, in the caspase-3 conformational landscape, the selection of P4 aspartate *versus* P4 valine is due to the combination of hydrogen bonding in the S4 pocket coupled to the network of hydrogen bonds that stabilize CP-R161. Changing the hydrogen bonding network of CP-R161 results in relaxed specificity in the case of DrCasp-3a. Given the constraints in the conformational landscape, it appears that multiple mutations are required to de-evolve the enzyme sufficiently in order to swap enzyme specificity. In contrast, changes in hydrogen bonding networks, within the same conformational landscape, are sufficient to improve activity against the second substrate, VEID, without significantly altering activity against the first substrate, DEVD. More generally, if cells do not require the separation of substrate selection, then fewer substitutions are required to evolve a dual function protease, as in caspase-3a, compared to neofunctionalization of enzyme specificity, as in the cases of caspase-3 *versus* caspase-6.

## Materials and Methods

### Cloning, expression, and purification

Zebrafish caspase-3a CP-N163T and CP-N163V were made by PCR site-directed mutagenesis and were cloned into pET11a expression plasmid containing either wild-type caspase-3 or zebrafish caspase-3a with a C-terminal histidine tag, as described previously (9, 17). The remaining mutants were made by GenScript. Mutations were confirmed by Sanger sequencing of both DNA strands. Plasmids were transformed into *E. coli* BL21(DE3) pLysS, and proteins were expressed as described (17, 18).

Caspase-3, −6 and −7 sequences in all species were downloaded from the CaspBase (13). A multiple sequence alignment (MSA) was generated for each subfamily by analyzing the amino acid frequency at each position with ProtParam on the ExPASy server (19). Sequence logos were generated utilizing the web-logo server (20).

### Enzyme activity assays

The enzymatic activity of the caspases were measured at 25 ℃ in a buffer of pH 7.5, 150 mM Tris, 50 mM NaCl, 0.1% sucrose, 0.1% CHAPS and 10 mM DTT, as described previously (21). Briefly, the total reaction volume was 200 μL, with a final enzyme concentration of 10 nM. The concentration of substrates Ac-DEVD-AFC or Ac-VEID-AFC were varied, as described previously (14). Samples were excited at 400nm, and the emission was monitored at 505 nm for 60 seconds. The steady-state parameters, K_M_ and k_cat_, were determined from plots of initial velocity *versus* substrate concentration.

### Phage display substrate libraries and selection

Caspase substrate libraries were constructed with an amino-terminal hexa-histidine sequence rendering the phage capable of binding to HisPur Ni-NTA resin, and caspase selection was determined as described previously (9). Briefly, phage libraries consisting of caspase cleavage sequences were bound to Ni-NTA resin, and the column was washed with a buffer of 50 mM Imidazole, 1 M NaCl, 1×PBS and 0.1% Tween-80 to remove unbound phage. Caspase enzyme (10-100 nM) was added to initiate the reaction, and samples were incubated for 3 hours. Phage possessing a suitable caspase cleavage site were released into the solution supernatant, and were amplified in *E. coli* ER2738 cells. The resulting phage were used for the following round of selection. Substrate sequences were determined after 3-5 rounds of selection, and the endpoint of the experiment was determined by plaque counting, where the number of phage bound to the resin was similar to the number of phage released during the treatment. The data showed no difference between the third, fourth, and fifth rounds, so data were combined.

### Crystallization data collection and molecular dynamics simulations

DrCasp-3a(CP-N163T) was crystallized as described previously for HsCasp-3 (14). Briefly, protein was dialyzed in a buffer of 10 mM Tris-HCl, pH 8.5, 1 mM DTT, concentrated to 4 mg/mL, and inhibitor Ac-DEVD-CHO (reconstituted in DMSO) was added at a 5:1 (w/w) inhibitor/protein ratio. Crystals were obtained at 18 ℃ by the hanging drop vapor diffusion method using a 4 μL drop that contained equal amounts of protein and reservoir solution. Each well contained a reservoir solution (500 μL) of 100 mM sodium citrate, pH 5.4, 23% PEG 6000, 10 mM DTT, and 3 mM NaN3. Flat, sheet-like crystals appeared within 14 days, and we used microseeding to obtain diffraction-quality crystals. In this case, crystal trays were set up as described above and incubated for 24 hours. The flat sheet crystals were collected and treated with seed beads using a kit from Hampton Research. Briefly, a 4 μL drop containing flat sheet crystals was added to a tube containing seed beads, following the manufacturer’s instructions. Reservoir solution (10 μL) was pipetted on the cover slide to remove all crystals from the coverslip, and the procedure was repeated five times. The resulting mixture of crystals and seed beads was vortexed for 30 seconds and cooled on ice for 10 seconds, and the procedure was repeated six times. Serial dilutions of the treated crystals were set up from 10^−1^ to 10^−3^, and 0.5 μL of the 10^−2^ dilution crystal seeds was added into the drops of the 24-hour crystal tray. Larger cube-shaped crystals appeared within 14 days. The crystals were collected and frozen in liquid nitrogen following the addition of 20% MPD (2-methlpentane-2,4-diol) plus the reservoir solution. Data were collected at 100 K at the SER-CAT synchrotron beamline (Advance Photon Source, Argone National Laboratory Argonne, IL, U.S.A). The data set contains 180 frames at 1°rotation. The protein crystallized in the orthorhombic space group P212121 and was phased with a previously published DrCasp-3a (PDB ID 5JFT). Data reduction and model refinements were done using HKL2000, COOT and Phenix (22–24).

Molecular dynamics (MD) simulations were performed using GROMACS 2016, as described previously (25–28). Simulations used the Amber99 force field and the TIP3P water model. Simulations started with the structure from wild-type DrCasp-3a (9), and mutations were added using Pymol. The proteins were solvated in a periodic box of 62 X 48 X 66 A, with ~14,000 water molecules, as described for HsCasp3a (29). Sodium or chloride atoms were added to neutralize the charge on the system. Steepest descent was used to minimize the system, and the waters were then relaxed during a 20 ps MD simulation with positional restraints on the protein. Simulations (50 ns) were then run under constant temperature (300K) and pressure, and a time step of 2 fs used. Coordinates were saved every 5 ps, and the protein was equilibrated within 500 ps.

## Data deposition

The crystal structure of DrCasp-3a(CP-N163T) has been deposited in the Protein Data Bank, www.wwpdb.org under PDB ID code 7JL7.

## ACKNOWLEDGEMENT

This work was supported by a grant from the National Institutes of Health [grant number GM127654 (to A.C.C)] and by funds from UT Arlington [Office of the Vice President for Research (to A.C.C)]. Use of the Advanced Photon Source was supported by the U.S. Department of Energy, Office of Science, Office of Basic Energy Sciences, under contract number W-31-109-ENG-38

