## Supplemental Figures Tables for "Remodeling hydrogen bond interactions results in relaxed specificity of Caspase-3"

Running title: Active site mutations relax specificity of Caspase-3

**Key Words**: enzyme specificity; apoptosis; caspase; zebrafish; protein evolution

### Supplemental Figures

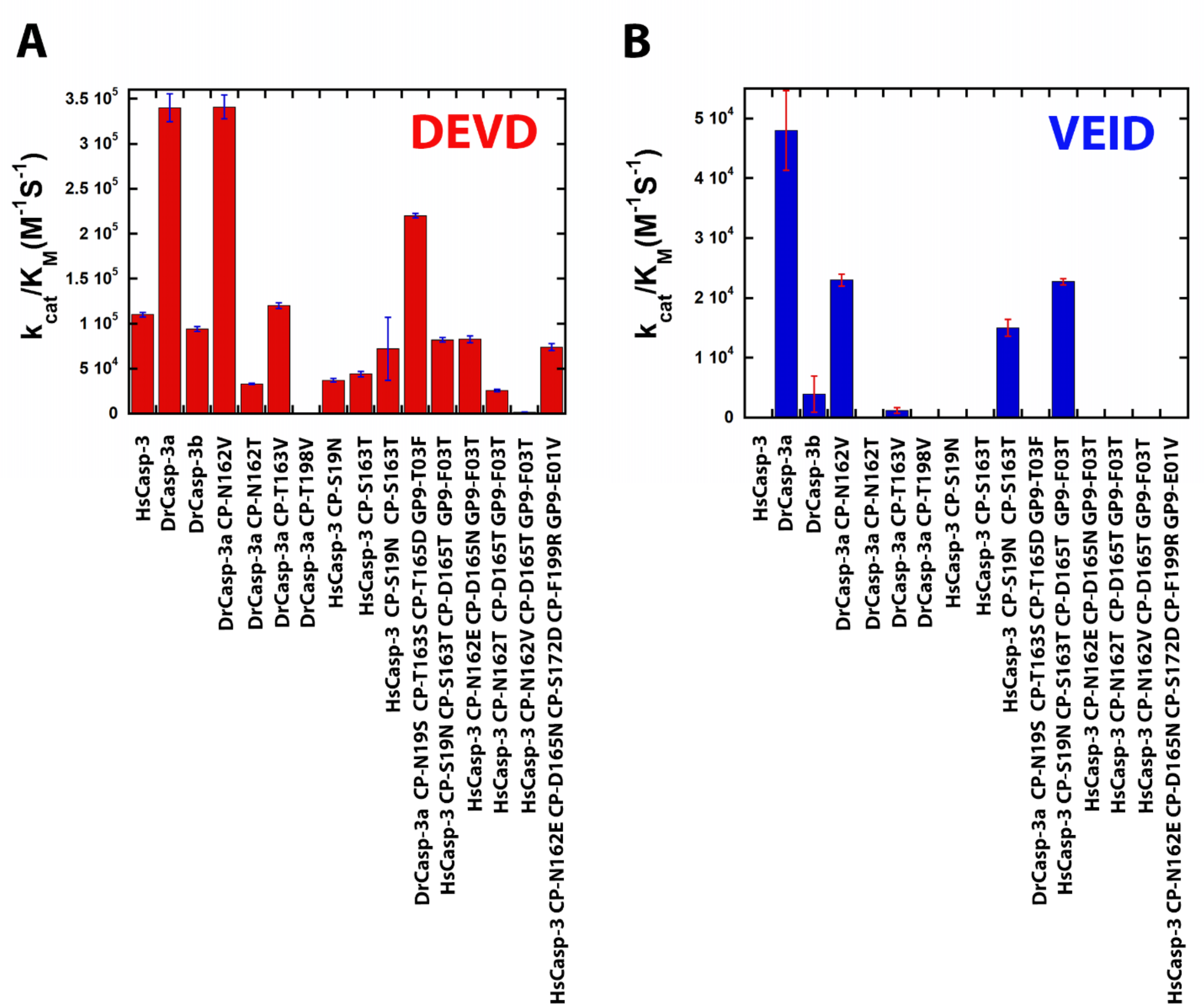


**Supplemental Figure 1. Enzymatic activity of caspase mutants.** Panels A,B. Enzyme specificity ($k_{cat}/K_M$) for wild-type and mutant caspases with DEVD (panel A) or VEID (panel B) as substrate. Data correspond to Table 2 in the main text.

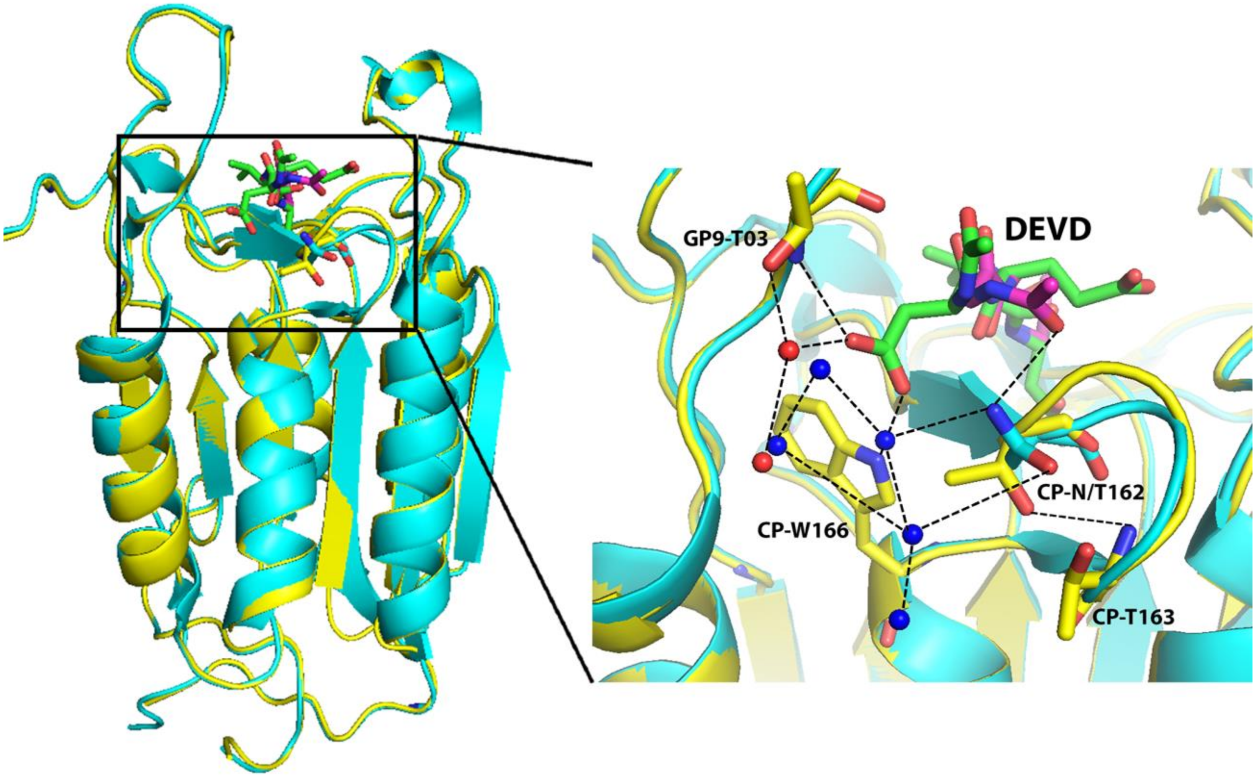


**Supplemental Figure 2. Crystal structure of DrCasp-3a CP-N162T.** Wild-type DrCasp-3a is shown in cyan, DrCasp-3a(CP-N162T) in yellow (PDB ID 7JL7). The inhibitor, DEVD, is shown in magenta for wild-type DrCasp3a and in green for DrCasp3a(CP-N162T). Water molecules are shown in blue for wild-type DrCasp3a and in red for DrCasp3a (CP-N162T).

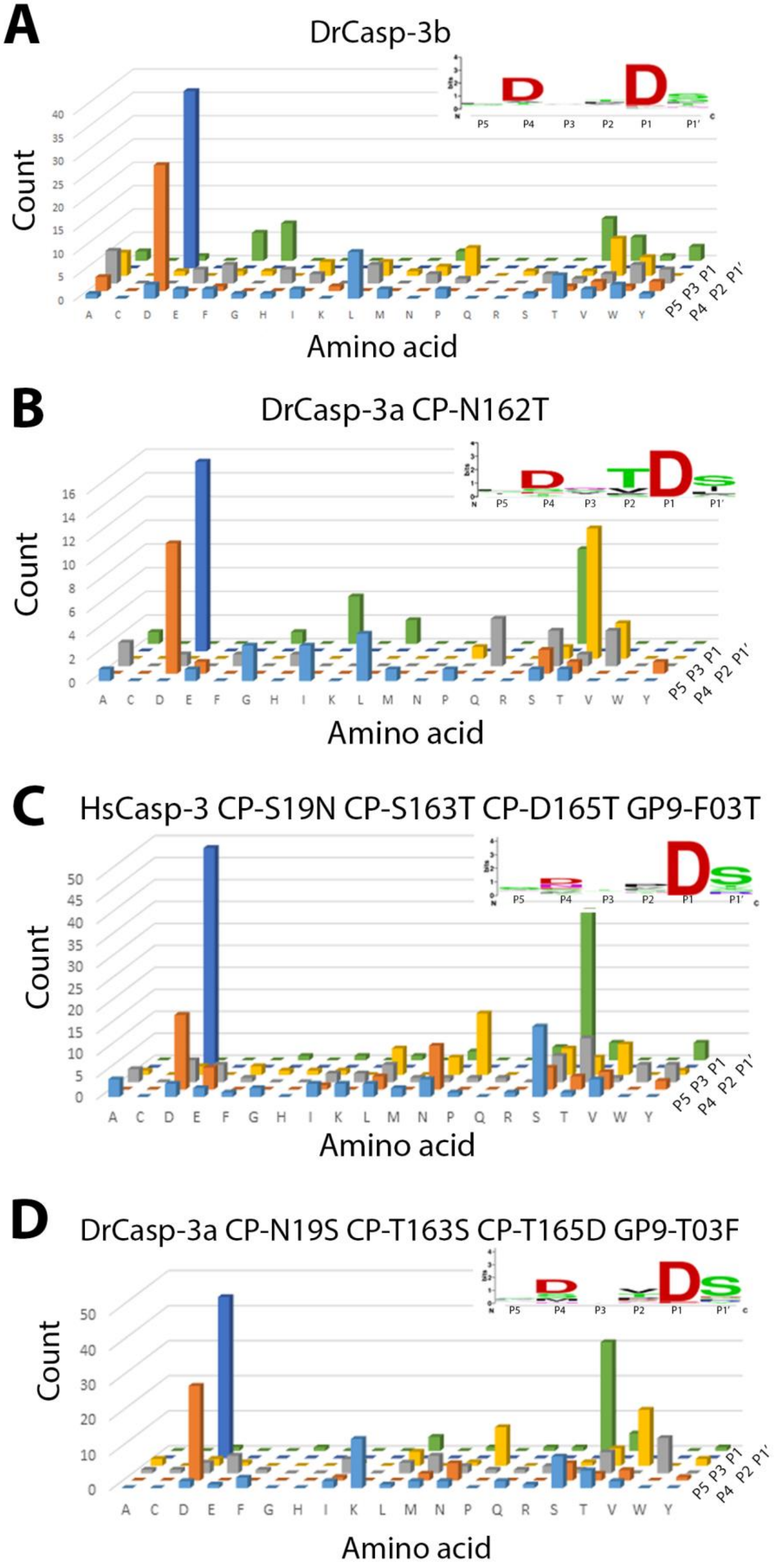
A
DrCasp-3b
Count
0 5 10 15 20 25 30 35 40
A C D E F G H I K L M N P Q R S T V W Y
P5 P3 P1
P4 P2 P1'
Amino acid
B
DrCasp-3a CP-N162T
Count
0 2 4 6 8 10 12 14 16
A C D E F G H I K L M N P Q R S T V W Y
P5 P3 P1
P4 P2 P1'
Amino acid
C
HsCasp-3 CP-S19N CP-S163T CP-D165T GP9-F03T
Count
0 5 10 15 20 25 30 35 40 45 50
A C D E F G H I K L M N P Q R S T V W Y
P5 P3 P1
P4 P2 P1'
Amino acid
D
DrCasp-3a CP-N19S CP-T163S CP-T165D GP9-T03F
Count
0 10 20 30 40 50
A C D E F G H I K L M N P Q R S T V W Y
P5 P3 P1
P4 P2 P1'
Amino acid

**Supplemental Figure 3. Substrate specificity determined by substrate phage display.** A-D. Substrate phage display for DrCasp-3b and mutants of HsCasp-3 and of DrCasp-3a. Sequence preferences are shown in the sequence logo inset for each panel. Results confirm DrCasp-3a(CP-N162T) and DrCasp3a(CP-N19S CP-T163S CP-T165D GP9-T03F) have high selection for DxxD, while HsCasp-3(CP-S19N CP-S163T CP-D165T GP9-F03T) have less selection for DxxD.
